# A superoxide reductase contributes to *Clostridioides difficile* resistance to oxygen

**DOI:** 10.1101/2022.12.19.521142

**Authors:** Rebecca Kochanowsky, Katelyn Carothers, Bryan Angelo P. Roxas, Farhan Anwar, V.K. Viswanathan, Gayatri Vedantam

**Affiliations:** School of Animal and Comparative Biomedical Sciences, The University of Arizona, Tucson, AZ, USA; BIO5 Institute for Collaborative Research, The University of Arizona, Tucson, AZ, USA; Southern Arizona VA Healthcare System, Tucson, AZ

**Author notes:** Correspondence to: Gayatri Vedantam, Ph.D., University of Arizona, School of Animal & Comparative Biomedical Sciences, 1117, E. Lowell St. Bldg. 90, Room, Tucson, AZ 85721. **Author Contributions** RK and KC generated the various strains, performed all experiments related to SOR activity and oxygen sensitivity and compiled the manuscript. FA provided technical support for the toxin and sporulation assays. BPR generated and analyzed the proteomics data and facilitated the structural comparisons. GV and VKV conceptualized and funded the studies, finalized the manuscript, and provided project oversight. RK and KC contributed equally to this study. VKV and GV contributed equally as senior authors. All authors read and approved the manuscript.

**Keywords:** *Clostridioides difficile*, superoxide reductase, redox, rubredoxin oxidoreductase

## Abstract

*Clostridioides difficile* causes a serious diarrheal disease and is a common healthcare-associated bacterial pathogen. Although it has a major impact on human health, mechanistic details of *C. difficile* intestinal colonization remain undefined. *C. difficile* is highly sensitive to oxygen and requires anaerobic conditions for *in vitro* growth. However, the mammalian gut is not entirely devoid of oxygen, and *C. difficile* tolerates moderate oxidative stress *in vivo*. The *C. difficile* genome encodes several antioxidant proteins, including a predicted superoxide reductase (SOR) that is upregulated upon exposure to antimicrobial peptides. The goal of this study was to establish SOR enzymatic activity and assess its role in protecting *C. difficile* against oxygen exposure. Insertional inactivation of *sor* rendered *C. difficile* more sensitive to superoxide indicating that SOR contributes to antioxidant defense. Heterologous *C. difficile sor* expression in *Escherichia coli* conferred protection against superoxide-dependent growth inhibition, and the corresponding cell lysates showed superoxide scavenging activity. Finally, a *C. difficile* SOR mutant exhibited global proteome changes under oxygen stress when compared to its parent strain. Collectively, our data establish the enzymatic activity of *C. difficile* SOR, confirm its role in protection against oxidative stress, and demonstrate its broader impacts on the vegetative cell proteome.

**Importance:** *Clotridioides difficile* is an important pathogen strongly associated with healthcare settings and capable of causing severe diarrheal disease. While considered a strict anaerobe *in vitro, C. difficile* has been shown to tolerate low levels of oxygen in its mammalian host. Among other well-characterized antioxidant proteins, the *C. difficile* genome includes a predicted superoxide reductase (SOR), an understudied component of antioxidant defense in pathogens. The significance of the research reported herein is the characterization of the enzymatic activity of the putative SOR protein, including confirmation of its role in protection of *C. difficile* against oxidative stress. This furthers our understanding of *C. difficile* pathogenesis and presents a potential new avenue for targeted therapies.

## Introduction

*Clostridioides difficile*, a Gram-positive spore-forming bacterium, is a major cause of antibiotic-associated diarrhea, and is responsible for significant numbers of healthcare-associated infections in Europe, Asia and North America (1). Over 400,000 *C. difficile* infections (CDI), with an associated 30,000 deaths, occur annually in the USA alone; these impose a >$1 billion burden in additional healthcare costs (2-4).

CDI is an intestinal disease, and presents as diarrhea, colitis and, sometimes, small intestinal damage (5). *C. difficile* cycles between two distinct morphotypes: a dormant spore form that is highly resistant to chemical and physical insults, and metabolically active vegetative cells that produce the toxins responsible for disease symptoms. Antibiotic suppression of the gut flora facilitates *C. difficile* spore germination and subsequent vegetative cell colonization. Toxigenic (toxin-producing) *C. difficile* strains harbor a 19.6kb genomic island (Pathogenicity Locus; PaLoc), that encodes the toxins TcdA and TcdB. These toxins glucosylate host Rho GTPases and cause intestinal damage and pathology. Some strains also express an ADP ribosylase toxin known as Binary toxin (or CDT) (6).

*C. difficile* vegetative cells are considered fastidiously anaerobic, and are more sensitive to oxygen than many enteric bacteria (7, 8). Nevertheless, vegetative cells are exposed to, and can tolerate, oxygen in the mammalian digestive tract. Oxygen content in the gut decreases progressively from the stomach to the small intestine and the colon (9-12); there is also an oxygen gradient from the central portion of the lumen, which is essentially anaerobic (pO_2_<1 mmHg), towards the epithelial surface, where oxygen concentrations approach 40mm Hg due to perfusion from underlying capillaries (12-14). Oxygen levels are also elevated in the dysbiotic gut, and vegetative cells have been observed in close proximity to the relatively oxygen-rich mucus layer (15-18). Correspondingly, *C. difficile* vegetative cells tolerate low levels of oxygen, displaying no loss of viability after 15 minutes of air exposure (7). Vegetative cells can grow at lower oxygen tensions in the range of 1-3% (corresponding to 7.6 – 22.8 mmHg) (19), and can also persist in ambient oxygen conditions in human intestinal organoids (20).

Recent studies have implicated a role for various *C. difficile* genes in resistance to reactive oxygen species (21-24) (Supplementary Table S1). A three-gene operon (*rbr1-perR-sor*) encoding a rubrerythrin, the transcriptional regulator PerR, and a putative superoxide reductase (SOR; desulfoferrodoxin) (Figure 1) was implicated in oxidative stress response in *C. difficile* (25). Specifically, a single amino acid change (T41A) in the peroxide-sensing transcriptional repressor PerR abrogated its ability to bind DNA, resulting in constitutive de-repression of the *rbr1-perR-sor* operon, and conferred increased oxygen tolerance to the laboratory-adapted strain 630Δ*erm* (25). This operon has canonical -10 and -35 sequences of a sigma factor A (σA)-dependent promoter (24), but its expression also appears to be under indirect control of the stress-response-related sigma factor σB (22). We previously observed increased abundance of the putative SOR in *C. difficile* exposed to the human antimicrobial peptide LL-37 (26), suggesting a potential role for this protein in intestinal colonization. Consistent with this, SOR expression was also upregulated in *C. difficile-*infected pig ileal loops (27).

**Figure 1.**
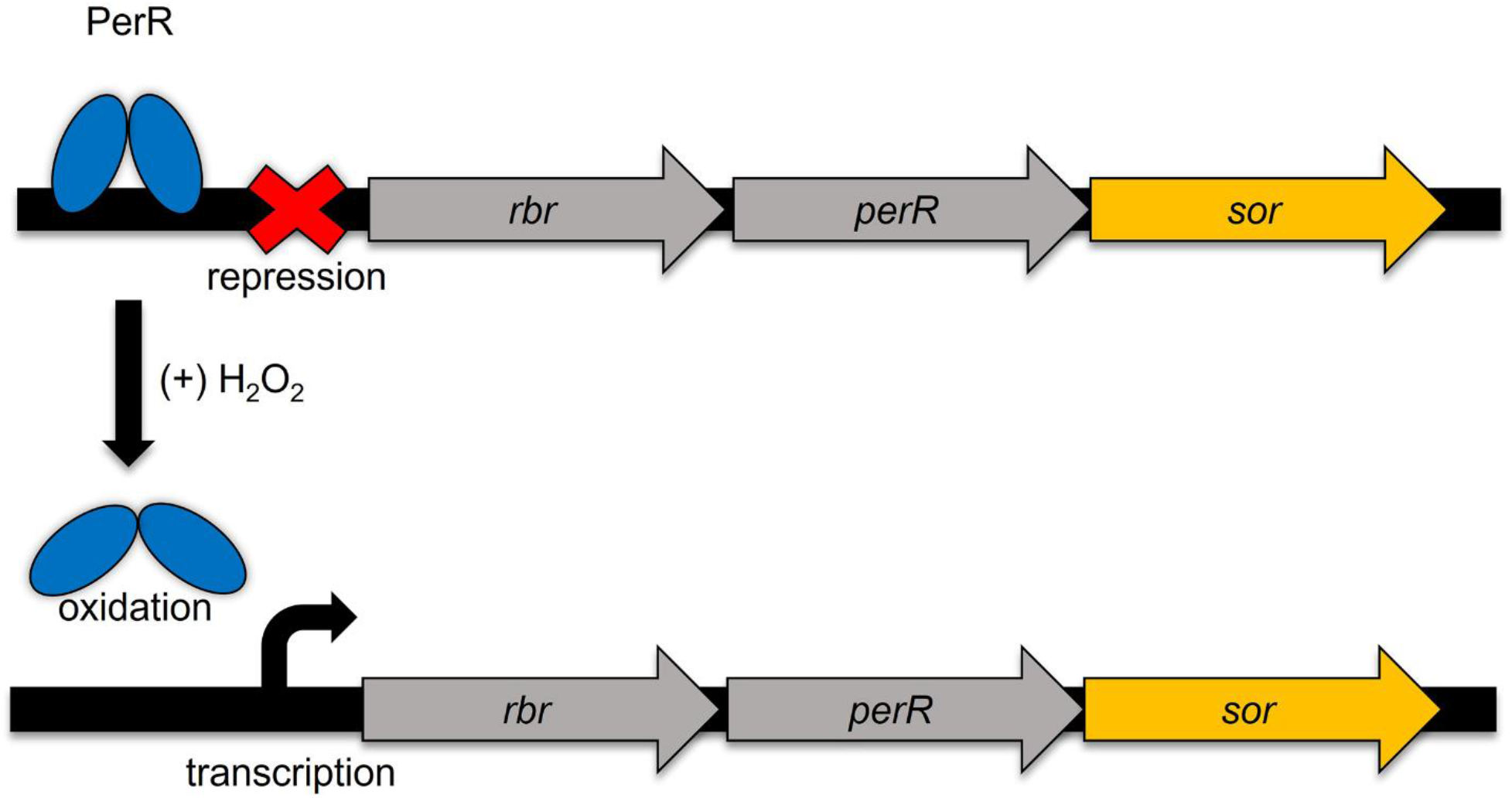
Genomic context of *C. difficile sor*. Diagram of the three-gene operon containing rubrerythrin (*rbr*), the PerR regulator (*perR*) and the putative superoxide reductase (*sor*). Oxidation by peroxide releases PerR from the operator site and derepresses the operon.

The precise mechanisms by which the proteins encoded by the *rbr1-perR-sor* operon contribute to oxidative stress resistance in *C. difficile* is not known. We generated a *sor* insertional mutant (*sor*::*erm*) and a corresponding plasmid-complemented strain (*sor*::*erm*/p*sor*) in the clinically-dominant ribotype 027 *C. difficile* strain BI-1 and compared their oxygen and reactive oxygen species (ROS) sensitivities to the isogenic parent strain. We also heterologously expressed *C. difficile sor* in an *Escherichia coli* strain lacking cytosolic superoxide dismutase (SOD), and assessed for ROS sensitivity, and for the ability of extracts to deplete superoxide. Finally, we compared the global protein expression profile of the *C. difficile sor*::*erm* mutant and the isogenic parent strain under oxidative stress conditions. Our studies confirm that *sor* expresses a functional superoxide reductase and protects *C. difficile* vegetative cells against oxidative stress.

## Results

### *C. difficile* genome encodes a superoxide reductase homolog

The superoxide anion (O_2_.^-^), formed by one-electron reduction of oxygen, and other reactive oxygen species (ROS) molecules derived from it are highly toxic to living systems. Superoxide reductases, and the better-studied superoxide dismutases (SODs), scavenge superoxide and mitigate this toxicity (28-30). SORs are phylogenetically diverse mononuclear iron enzymes that reduce superoxide to hydrogen peroxide (31, 32). The *C. difficile* genome encodes a 128 amino acid (aa) SOR homolog as the third and last gene (open reading frame CD0827) in an operon implicated in resistance to redox stress (Figure 1) (25). CD0827 is conserved in all publicly-available *C. difficile* genomes. SORs are classified on the basis of the number of iron centers, and the *C. difficile* sequence (Figure 2A) and structure (Figure 2B, 2D) closely resembles the 2Fe-SOR (or desulfoferrodoxins) of *Desulfovibrio desulfuricans* (43% amino acid identity), one of the earliest characterized SOR proteins (31). Desulfoferrodoxins were previously called rubredoxin oxidoreductases (Rbo). The FeCys_4_ Center I (Desulforedoxin, Dx) of 2Fe-SORs (Figure 2C) is dispensable for enzymatic activity; the superoxide scavenging Center II (Neelaredoxin, Nlr) has an iron coordinated by four histidines and one cysteine residue [Fe(His)4(Cys)] that are essential for activity, as well as a conserved glutamate (32, 33) (Figure 2C). Based on conservation of the essential active site residues in the CD0827-encoded amino acid sequence, and structural similarity to *D. desulfuricans* SOR we sought to confirm the enzymatic activity of the putative *C. difficile* SOR and assess its role in oxygen stress tolerance.

**Figure 2.**
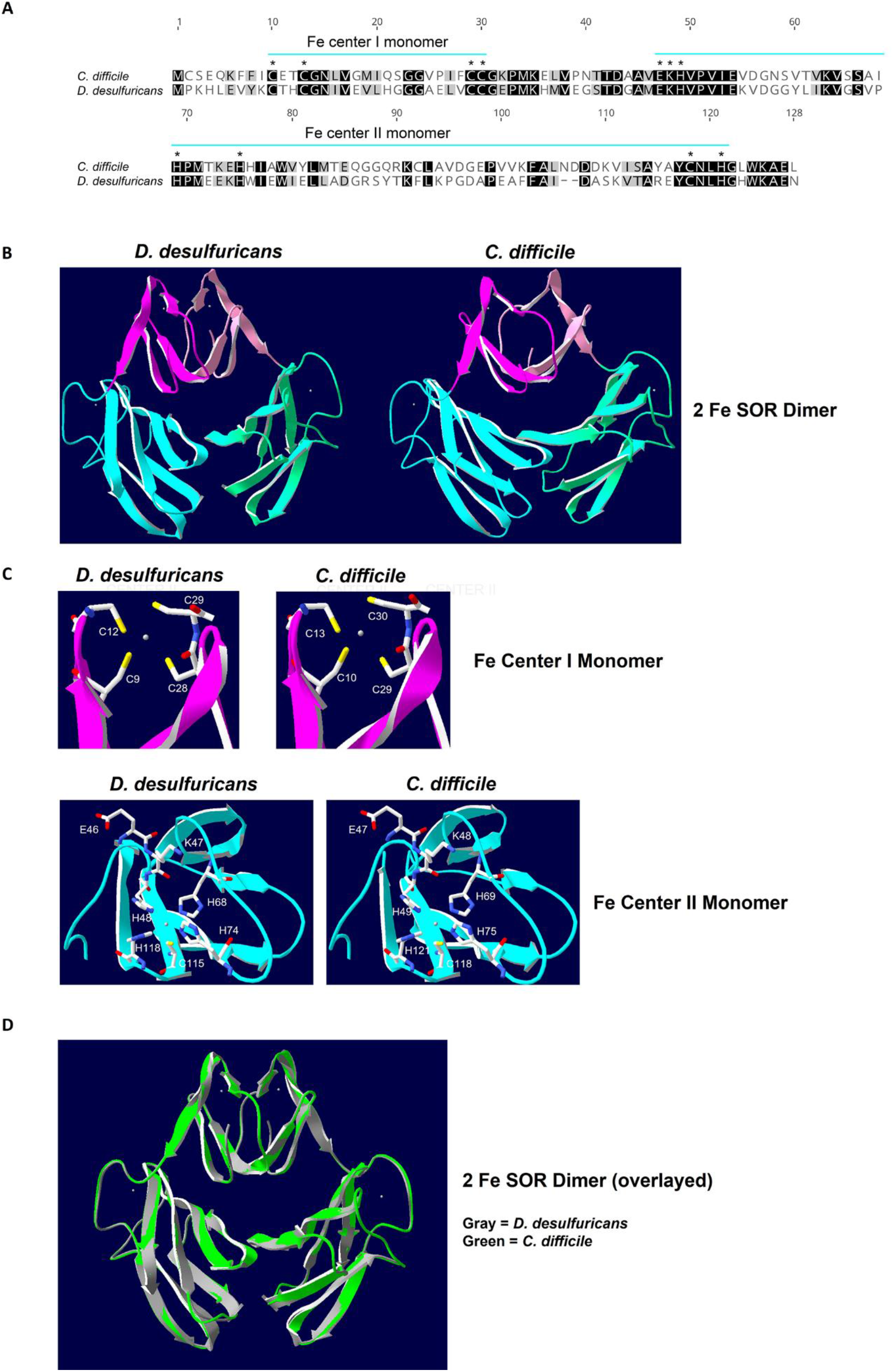
Structural conservation of *C. difficile* SOR. (A). Residues indicated in the black box are identical, while residues in the gray box are similar. * is used to identify the residues involved in Fe binding and blue lines are used to show the boundaries of the Fe centers. (B). Structures of superoxide reductase dimer in *D. desulfuricans* (left) and *C. difficile* (right). The FeCys_4_ Center I (Dx domain) of each monomer is shown in pink and the [Fe(His)4(Cys) Center II (Nlr domain) is shown in blue. (C). Structures of superoxide reductase Fe center I (pink) and II (blue) and conserved catalytic residues in *D. desulfuricans* (left) and *C. difficile* (right). Iron atoms are shown as gray spheres. (D). Structural alignment of superoxide reductase dimer in *D. desulfuricans* (gray) and *C. difficile* (green).

### SOR mitigates *C. difficile* sensitivity to air and ROS

To assess the contribution of *C. difficile sor* to protection against oxidative stress, we generated an insertional disruption of *sor* (*sor::erm*) via ClosTron mutagenesis (34). A corresponding plasmid-complemented strain (*sor::erm/*p*sor*) and a vector-only control strain were also generated. Wild type (WT) BI-1, *sor*::*erm* and *sor*::*erm*/p*sor* strains were exposed to ambient air for defined periods, and surviving bacteria were enumerated. Consistent with previous reports of *C. difficile* 630Δ*erm* (35), ∼50% of WT BI-1 was recovered after a 30-minute air exposure (Figure 3), and no viable bacteria were recovered after a 60-minute treatment (data not shown). In contrast, survival of the SOR-deficient *sor*::*erm* strain was significantly reduced after 30 minutes of air exposure compared to WT BI-1, with only ∼12% of the starting CFU recovered. Complementation with p*sor* restored viability to WT levels (Figure 3). This suggests that *sor* limits *C. difficile* sensitivity to oxygen.

**Figure 3.**
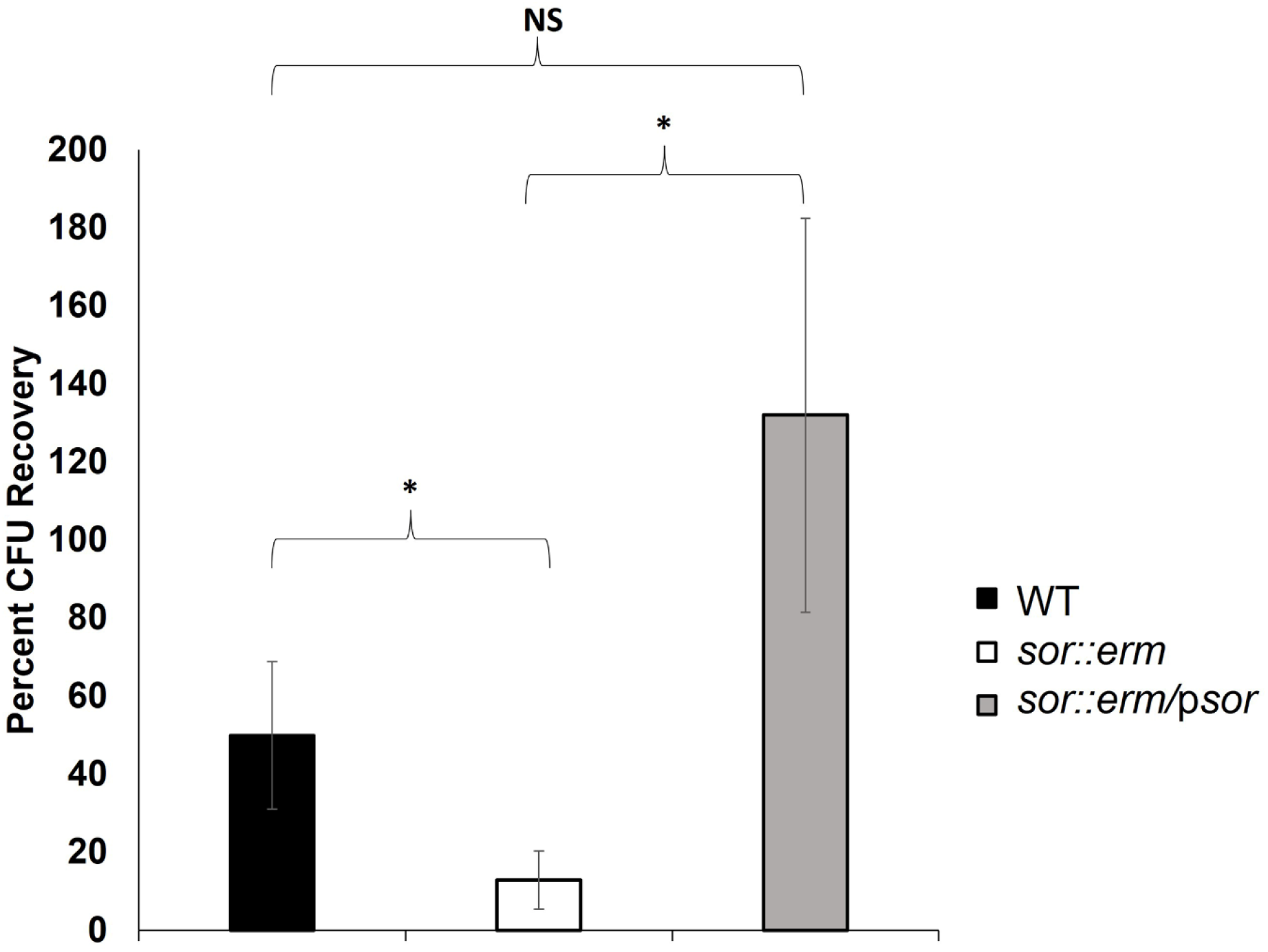
SOR disruption reduces *C. difficile* viability under oxygen stress. WT BI-1, *sor::erm*, and *sor::erm/*p*sor* cultures were exposed to ambient oxygen for 30 minutes and surviving CFUs were enumerated by plating. Viability under oxygen stress was calculated using CFU recovered as a percentage of starting CFUs. Cultures were compared in biological triplicate and means were tested for statistical significance by ANOVA. Tukey’s post-hoc test was used for pairwise comparisons. *p<0.05; NS = not significant. Error bars represent standard deviation.

To address whether *C. difficile* SOR specifically protects against superoxide, we assessed sensitivity to menadione (MD), which promotes the conversion of molecular oxygen to superoxide (Figure 4A). Under anaerobic conditions, WT, *sor::erm* and *sor::erm/*p*sor* displayed comparable menadione sensitivities. In contrast, air exposure increased *C. difficile* WT sensitivity to menadione, and this was significantly more pronounced for *sor::erm;* plasmid complementation reversed the phenotype (Figure 4B).

**Figure 4.**
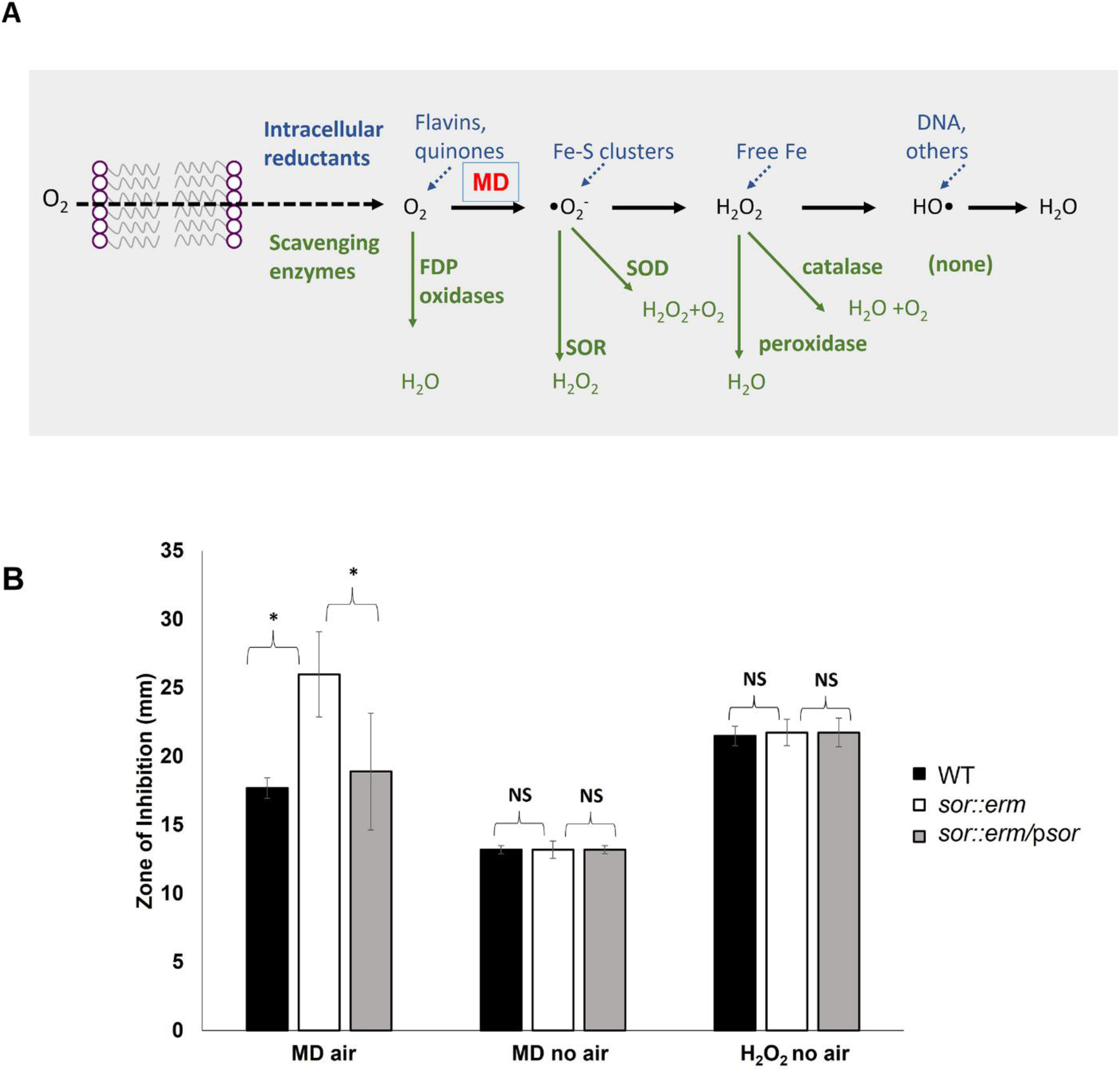
SOR protects *C. difficile* from ROS. (A). In aerobic conditions, menadione (MD) promotes the conversion of molecular oxygen to superoxide, which can be scavenged by SOD and SOR. Superoxide is not generated in anaerobic conditions. Multiple enzymes, including SOR, participate in the detoxification of molecular oxygen, reactive oxygen species, and peroxide. (B). Under aerobic conditions, SOR disruption reduced the tolerance to superoxide generated by menadione, as measured by zone of inhibition (ZOI). SOR did not impact *C. difficile* tolerance to H_2_O_2_. ZOIs were compared in triplicate. *p<0.05; NS = not significant. Error bars represent standard deviation.

While MD treatment augments intracellular superoxide abundance, it can also increase H_2_O_2_ and OH• levels (Figure 4A). To definitively establish a role for *C. difficile* SOR in protection against superoxide, we assessed sensitivity to H_2_O_2_ under anaerobic conditions (which will not increase superoxide levels; Figure 4A). WT, *sor::erm* and *sor::erm/*p*sor* did not exhibit significant differences in their susceptibility to H_2_O_2_ (Figure 4B). Together, our data suggest that *sor* promotes *C. difficile* aerotolerance by specifically protecting against superoxide.

### *C. difficile* SOR is a superoxide scavenger

To verify superoxide reductase activity, *C. difficile sor* was cloned into the pTrcHis2 TOPO vector, in-frame with the C-terminal *myc-6Xhis* tag sequence. The resulting plasmid, p*CDsor* was introduced into *Escherichia coli* QC779, which lacks two endogenous cytoplasmic SODs (SodA and SodB) and is, therefore, sensitive to superoxide-generating compounds like menadione (36). IPTG-inducible, heterologous SOR expression in this background was verified by immunoblotting against the Myc tag (Supplementary Figure S1). In zone of inhibition studies, the SOR-expressing *E. coli* strain (QC779/p*CDsor*) was significantly more resistant to MD compared to the parent strain, or a strain harboring the empty vector (QC779/vector; Figure 5A), confirming a role for C. *difficile* SOR in protection against superoxide.

**Figure 5.**
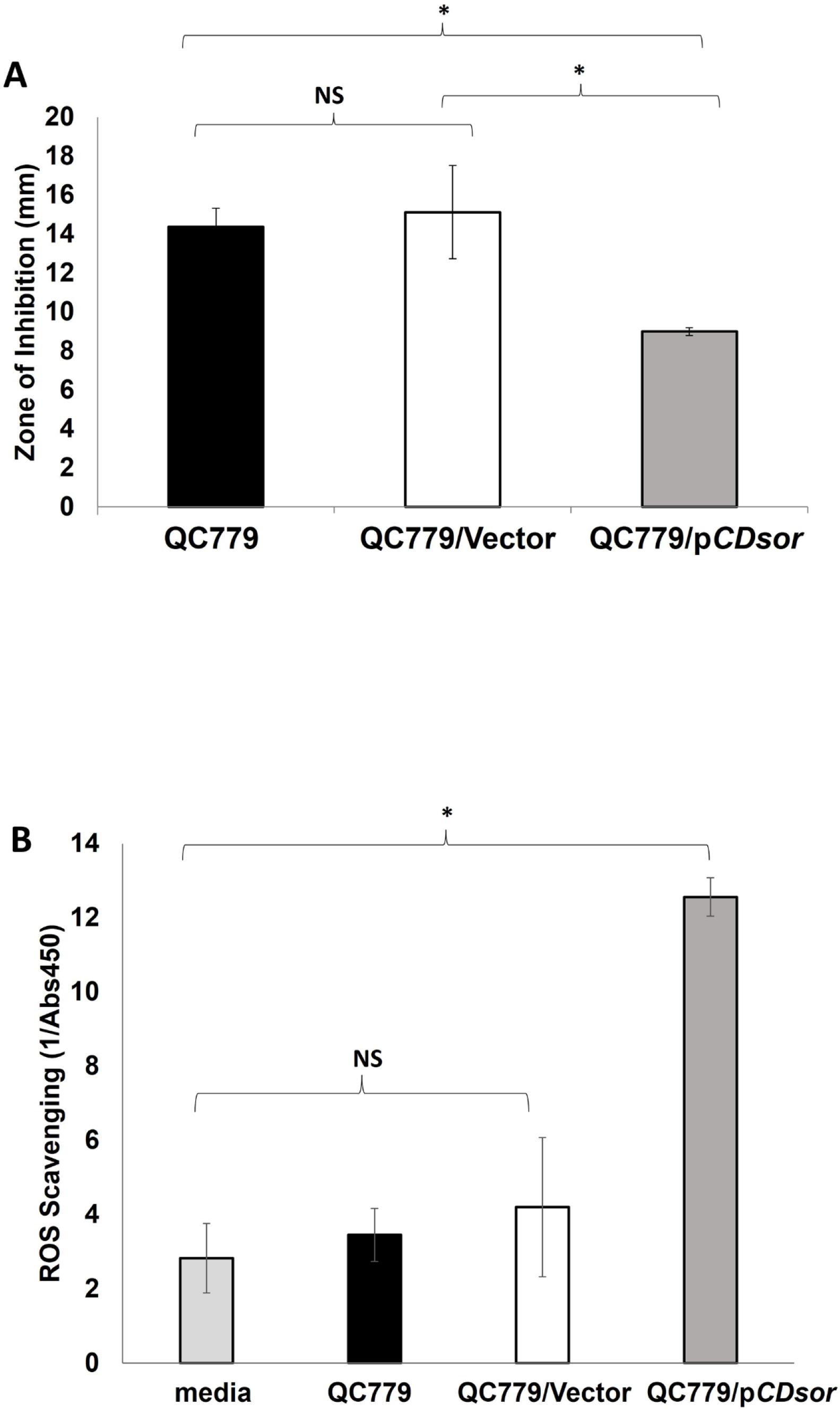
Expression of SOR in *E. coli* QC779 rescues ROS scavenging. (A). The menadione sensitivity of QC779 and its derivatives were measured by disc diffusion. (B). ROS scavenging capacity for lysates of QC779 and its derivatives was measured by a colorimetric change in substrate in the presence of xanthine oxidase after 1 hr. Experiments were conducted in biological triplicate. *p<0.05; NS = not significant. Error bars represent standard deviation.

To quantitatively assess ROS scavenging activity of QC779 and its derivatives, we utilized a xanthine oxidase (XO) assay. XO catalyzes the conversion of xanthine and oxygen into uric acid, hydrogen peroxide, and superoxide. In the XO assay, the superoxide anion converts a tetrazolium salt into a formazon dye; superoxide depletion, for instance by SOR or SOD, decreases dye formation. In this assay, QC779/p*CDsor* lysates displayed significantly higher superoxide scavenging capacity compared to WT QC779 and QC779/Vector in the presence of xanthine oxidase (Figure 5B).

Next, we used a standard in-gel assay to further confirm SOR-dependent superoxide scavenging (37, 38). Cell extracts separated by native polyacrylamide gel electrophoresis were exposed to riboflavin, which accelerates aerobic superoxide generation. Subsequent treatment with the superoxide-reactive dye nitroblue tetrazolium (NBT) results in uniform staining of the gel except in areas corresponding to the presence of superoxide scavenging enzymes (37, 38). In this assay, a zone of clearing was observed for QC779/p*CDsor* cell lysates that was not present in the lanes corresponding to QC779 and QC779/vector (Supplementary Figure S2). The presence of a protein corresponding to the zone of clearing with QC779/p*CDsor*, but not QC779 WT or empty vector lysate, was confirmed by immunoblotting against the histidine tag (Supplementary Figure S2). As expected, the fusion protein was more dispersed in the native PAGE (Supplementary Figure S2) compared with the staining observed with the SDS PAGE in Supplementary Figure S1.

Together, these data confirm a role for SOR in scavenging superoxide and protecting *C. difficile* against ROS sensitivity. The catalytic iron centers of SORs are oxidized during the conversion of superoxide to peroxide, requiring a separate electron donor to regenerate the reduced enzyme capable of further catalytic activity. This is typically mediated by small iron-containing proteins called rubredoxins. A putative rubrerythrin encoded upstream in the same operon as *sor* (*rbr*, Fig. 1) likely plays this reducing role for *C. difficile* SOR. Since *C. difficile* SOR, by itself, exhibits activity in the heterologous *E. coli* system, our data suggest that this enzyme, like others in the class (39, 40), is promiscuous in accepting electrons from non-native donors.

### *C. difficile* SOR disruption impacts proteome abundance in response to oxygen stress

To investigate the role of SOR in global gene regulation under oxidative stress, LC–MS/MS and label-free quantitative proteomic analyses were used to compare WT and *sor::erm* strains grown under anaerobic conditions and subsequent to oxygen exposure. In WT BI-1, a total of 153 proteins were altered in abundance upon oxygen exposure compared to anaerobic conditions (73 increased, 80 decreased), while in *sor*::*erm*, only 66 proteins showed any abundance changes (38 increased, 28 decreased) following air exposure (Table 1, Supplementary Table S2). Specifically, and upon oxygen exposure, wild-type BI-1 had increased abundance of proteins with oxidoreductase activity, and those associated with histidine metabolism, and folylpolyglutamate metabolism. In contrast, proteins involved in peptidoglycan- and glutathione metabolism were decreased (Figure 6A).

**Table 1.** Number of proteins differentially abundant between WT BI-1 and BI-1 *sor::erm* in anaerobic and aerobic conditions. LC–MS/MS and label-free quantitative proteomic analysis were used to compare WT and *sor::erm* strains in anaerobic conditions and following 30 minutes of oxygen exposure. A fold change greater than 2 was defined as the cutoff for differentially abundant proteins.

|  |  | Numerator |  |  |  |  |
| --- | --- | --- | --- | --- | --- | --- |
|  |  | Strain | WT |  | *sor::erm* |  |
|  |  | O<sub>2</sub> | - | + | - | + |
| Denominator | WT | - |  | 73↑, 80↓ | 75↑, 39↓ | 76↑, 54↓ |
|  |  | + | 80↑, 73↓ |  | 98↑, 95↓ | 119↑, 102↓ |
|  | *sor::erm* | - | 39↑, 75↓ | 95↑, 98↓ |  | 38↑, 28↓ |
|  |  | + | 54↑, 76↓ | 102↑, 119↓ | 28↑, 38↓ |  |

**Figure 6.**
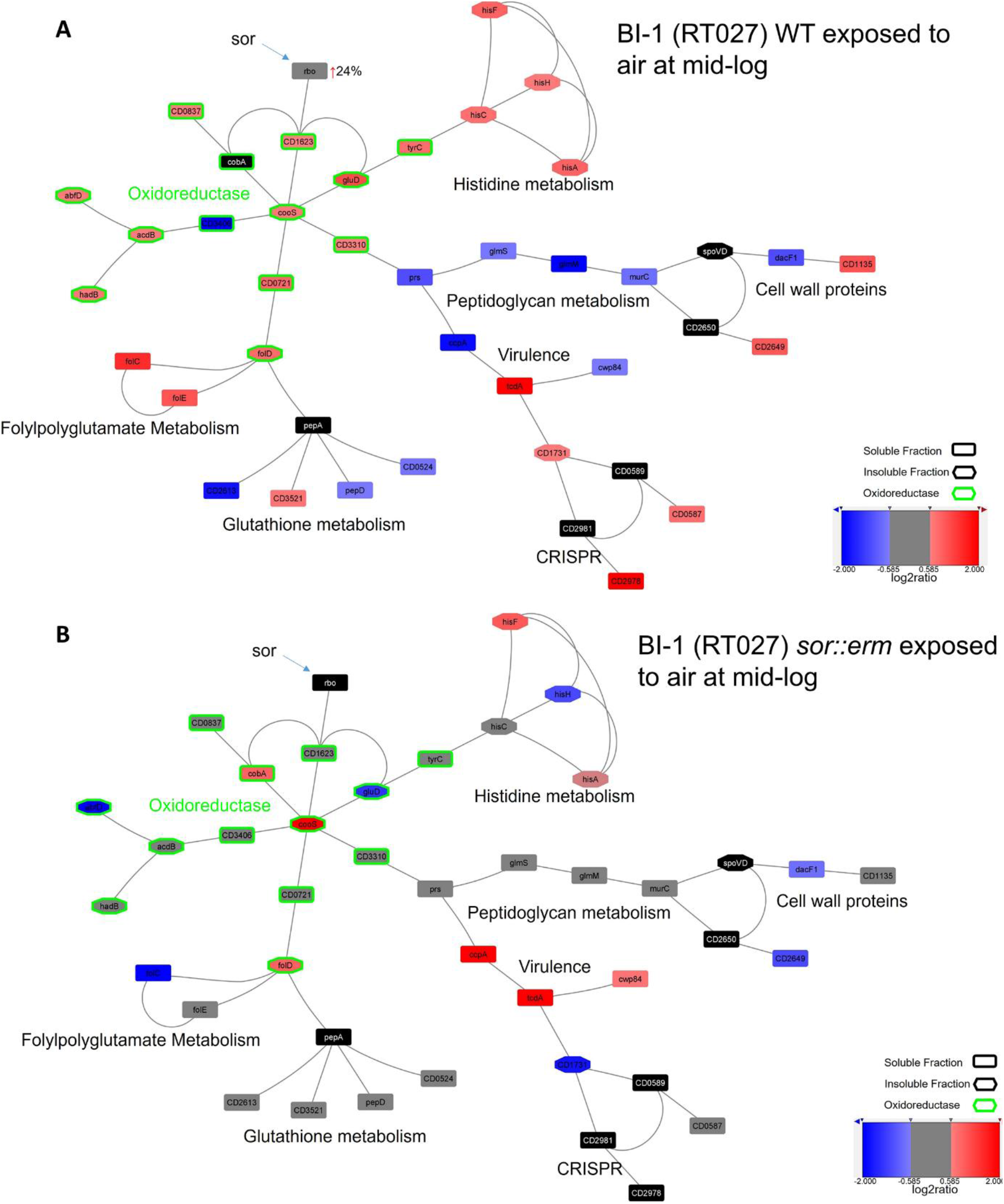
SOR disruption alters protein abundances in BI-1 vs BI-1 *sor::erm* following oxygen exposure. Protein-protein interaction (PPI) networks were constructed to compare BI-1 and *sor::erm* in the presence and absence of oxygen stress, using gene ontology (GO), Kyoto Encyclopedia of Genes and Genomes (KEGG), and Clusters of Orthologous Genes (COG) databases.

Strikingly, many fewer of the proteins above were altered in abundance during oxygen exposure if SOR was disrupted (Figure 6B). However, a total of 221 molecules were differentially regulated in *sor*::*erm* compared to the wild-type BI-1 strain following oxygen exposure (221 proteins; 119 upregulated, 102 downregulated). The large number of dysregulated proteins is likely due to the intracellular “high oxygen stress” environment present in this SOR-strain (Table 1, Supplementary Table S2). Curiously, even in the absence of air exposure, proteome abundance was altered in *sor*::*erm* compared to the isogenic parent (114 proteins; 39 increased, 75 decreased) suggestive of superoxide presence and/or a role for SOR even under anaerobic conditions. Our results suggest a differential *C. difficile* response to oxygen stress in the absence of SOR, and possible SOR-dependent regulation of multiple cellular processes, including those not known to be directly implicated in oxygen stress response.

### SOR disruption does not impact major *C. difficile* virulence traits

Given the extensive alterations in protein abundances, we explored if SOR/redox stress directly or indirectly influenced virulence-associated traits including toxin production, antibiotic resistance, and sporulation. Toxin production was not significantly altered between the BI-1 WT and the *sor::erm* derivative (Figure 7A), and *sor* disruption did not alter spore yields (Figure 7B). While sensitivity to a variety of antibiotics was not altered in *sor*::*erm* compared to the parent WT strain, both the WT and *sor::erm* derivatives displayed marked increases in sensitivity to metronidazole, cefoxitin and trimethoprim when combined with brief air exposure (Table 2).

**Table 2.** SOR disruption does not impact antibiotic resistance. Resistance to antibiotics was tested via minimum inhibitory concentration (MIC) and Zone of Inhibition (ZOI) assays. MICs were performed in triplicate.

| MIC (µg/mL) | WT | *sor::erm* | WT+air | *sor::erm*+air |
| --- | --- | --- | --- | --- |
| Metronidazole | 6.25 | 6.25 | 0.78 | 0.78 |
| Rifampicin | <0.05 | <0.05 | <0.05 | <0.05 |
| Cefoxitin sodium | 50 | 50 | 12.5 | 12.5 |
| Trimethoprim | 25 | 25 | 6.25 | 6.25 |
| ZOI (mm) | WT | *sor::erm* | WT+air | *sor::erm*+air |
| Penicillin | 0 | 0 | 0 | 0 |
| Streptomycin | 0 | 0 | 0 | 0 |
| Kanamycin | 0 | 0 | 0 | 0 |
| Chloramphenicol | 20.5 | 20 | 21 | 22 |
| Erythromycin | 19 | 0 | 21 | 0 |
| Moxifloxacin | 17 | 17 | 18 | 18 |

**Figure 7.**
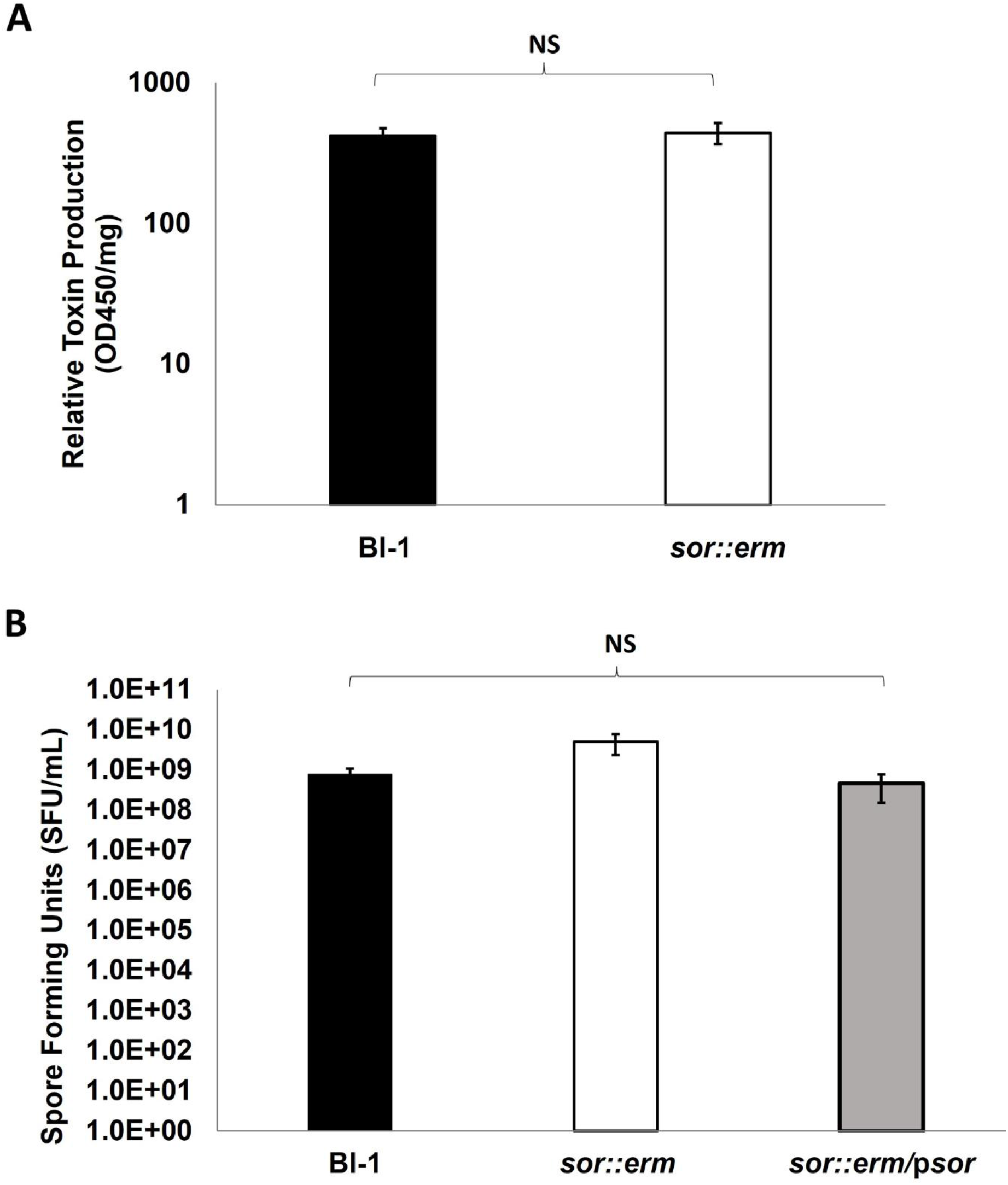
SOR disruption does not impact sporulation or toxin production. (A). Toxin production was measured by ELISA after 72 hours of anaerobic growth. (B). Spore production was measured after 12 days of growth, heat shock, and plating on BHI agar supplemented with taurocholate. Experiments were conducted in biological triplicate. NS = not significant. Error bars represent standard deviation.

## Discussion

The *C. difficile* genome encodes several proteins with roles in scavenging reactive oxygen species (ROS), the direct mediators of oxidative damage. Our studies demonstrate that *C. difficile* SOR is a functional superoxide scavenger and that it robustly mitigates redox stress. SORs are unusual enzymes, encoded in the genomes of only a few hundred sequenced bacteria and archaea, mostly anaerobes. Because the single predicted SOD of *C. difficile* is restricted to the spore (41), SOR appears to be the primary superoxide scavenger in the vegetative cells. Although the role of SORs in antioxidant defense has been established for several organisms (42-45), very few reports focus on the SORs of pathogens, and little is known about their role in infection. Giacani et al. showed that the *Treponema pallidum* SOR expression is increased during rabbit infection, under the control of an extra-cytoplasmic sigma factor (46).

SODs, on the other hand, are required for full virulence in animal models of *Campylobacter coli* (47), *Yersinia enterocolitica* (48), *Bordetella pertussis* (49), *Neisseria meningiditis* (50), *Streptococcus agalactiae* (51), and *Streptococcus pneumonia* (52), among others. In pathogens with intracellular phases, the attenuation of SOD mutants is often correlated with increased sensitivity to phagocyte killing; this has been shown in *Shigella flexneri, Bordetella pertussis* and *Salmonella typhimurium* (50). A *sodC* mutant in *Salmonella typhimurium* had attenuated virulence in a murine C57BL/6 model, but strikingly, this SOD mutant killed phox-91 knockout mice, which are deficient in the neutrophil oxidative burst, as effectively as wild type bacteria, specifically linking the attenuation of the SOD mutants in wild type animals to their defect in resisting phagocyte-derived ROS (53).

Overall, the preference of *C. difficile* for anaerobic growth is likely coupled with expression of various molecules that contribute to defense against oxidants (21–25, 41, 54) (Table 1). Permpoonpattana et al. demonstrated that a catalase, a peroxidase, and *C. difficile*’s single predicted superoxide dismutase (41) are present on the spore coat. The stress-response-related sigma factor σB was additionally shown to be essential for *C. difficile* tolerance to low oxygen concentrations. Mutants lacking *sigB* failed to grow in the presence of 0.1% oxygen, and exhibited a colonization defect in axenic mice (22). Four genes directly and positively regulated by σB have also been implicated in oxygen detoxification (24). These include the flavodiiron proteins FdpA and FdpF and the reverse rubrerythrins, revRbr1 and revRbr2. FdpF, revRbr1 and revRbr2 exhibit NADH-linked oxygen- and H_2_O_2_-reductase activities, while FdpA is primarily associated with an oxygen-reductase activity (23, 24). A mutant lacking all four proteins showed a growth defect at an oxygen concentration of 0.1%, and failed to grow in the presence of 0.4% oxygen (24).

Giordano and colleagues demonstrated a protective effect of a *C. difficile* cysteine desulfurase in low oxygen environments. Deletion of *iscS2*, a gene encoding a cysteine desulfurase likely involved in iron-sulfur cluster assembly, resulted in a severe growth defect in the presence of 2% oxygen (21). The mutant was also impaired for colonization in a mouse model of CDI.

In a study by Troitzsch et al, a spontaneous point mutation in the transcriptional repressor PerR in *C. difficile* 630Δ*erm* was shown to result in constitutive expression of oxygen stress-related proteins and higher oxygen tolerance (25). Notably, two groups have reported increased expression of putative ROS scavengers in *C. difficile* during animal infection. Janoir et al. observed increased transcription of rubrerythrin (*rbr*) during infection of germ-free mice (55), and Scaria et al. showed increases in rubrerythrin and desulfoferrodoxin (*sor*) gene expression in porcine ileal loops (27). We previously demonstrated SOR over-expression following *C. difficile* exposure to the human antimicrobial peptide LL-37, suggesting a role for SOR during infection (26). Our proteomics results also suggest a role for SOR in the regulation of multiple cellular processes, particularly during oxygen exposure, including redox stress and multiple forms of metabolism (Figure 6). Protein expression was altered to a lesser extent between the WT and *sor::erm* in anaerobic conditions, though a role for SOR beyond the oxygen stress response remains to be investigated.

Interestingly, we did not observe an impact of SOR disruption on major *C. difficile* virulence traits, including toxin production and sporulation, *in vitro*. Our proteomics data showed increased abundance of Toxin A following short-term oxygen exposure in both wild-type and *sor::erm* strains (Figure 6), though toxin production at 72 hours of anaerobic incubation was not impacted in a SOR-dependent manner (Figure 7). Sensitivity to multiple antibiotics was not altered due to SOR disruption, but sensitivity to metronidazole, cefoxitin and trimethoprim was increased in both strains following oxygen exposure. Oxygenation has been shown to enhance the efficacy of the antibiotic amikacin in killing *Mycobacterium abscessus* (56), and in *C. difficile*, resistance to metronidazole has been linked to mutations in the pyruvate-ferredoxin/flavodoxin oxidoreductase gene (*nifJ*) (57), though the link between oxygen stress response and antibiotic sensitivity is not well understood.

Taken together, our findings are consistent with a large body of literature describing the need for pathogens to combat superoxide stress during infection. Apart from extracellular sources, bacteria must also contend with ROS generated inside the cell when oxygen is present. Molecular oxygen easily crosses biological membranes and produces ROS by reacting with various biomolecules inside the cell (Figure 4A). Because superoxide is not membrane permeable (58), the main source of intracellular superoxide is thought to be this type of endogenous generation (59). Thus, expression of a bona fide superoxide scavenger suggests that *C. difficile* encounters substantial molecular oxygen—the source of superoxide—during the course of infection. This has important implications for our understanding of *C. difficile* biology, particularly because the organism is strictly anaerobic *in vitro*. Recent work has dispelled the notion that the gut is entirely anoxic. Marteyn et al. demonstrated the existence of a relatively oxygenated zone in the intestinal lumen close to the epithelial surface, and showed its impact on *Shigella flexneri* pathogenesis (60). Interestingly, Koenigsknecht et al. demonstrated the presence of *C. difficile* throughout the small and large intestines of mice (61), indicating that *C. difficile* may not be restricted to the most anaerobic portions of the GI tract as previously hypothesized. SOR-deficient mutants could be attenuated in virulence due to decreased survival in more oxygenated parts of the gut, a hypothesis our group is actively investigating.

Because we have verified the action of SOR on superoxide specifically via heterologous expression (Figure 5), we propose that SOR protects superoxide-sensitive components of *C. difficile* required during infection. These components present attractive drug targets. Superoxide is known to damage Fe-S clusters, present in many enzymes involved in anaerobic metabolism (59), and a number of these enzymes are increased in expression in models of *C. difficile* infection (27, 55). One such protein, PFOR, is already being exploited as the target of the antibiotics nitazoxanide and Amixicile (62, 63). PFOR is found mostly in anaerobes, and is therefore a narrow-spectrum anti-infective target. At least one study suggests that Amixicile is less disruptive to the gut microbiota and may decrease the risk of recurrent CDI compared to less targeted therapies (63). Other superoxide-labile *C. difficile* enzymes could also be exploited similarly.

In summary, we have shown that the superoxide scavenger SOR contributes to the survival and protein regulation of *C. difficile*, highlighting a potential vulnerability of this pathogen that should be considered in designing new therapies.

## Methods

### Bacterial strains and culture

*C. difficile* BI-1 was a kind gift from Dr. Dale Gerding. All *C. difficile* strains were routinely cultured anaerobically at 37°C. Because of their increased sensitivity to oxygen, SOR mutants were propagated on thioglycollate (TG) agar (Difco fluid glycollate medium with agar added to 1.6%). For assays comparing WT, *sor*::*erm* and *sor*::*erm*/p*sor*, all strains were grown on TG agar, supplemented with antibiotics when appropriate. For broth cultures, *C. difficile* strains were cultured in Brain Heart Infusion media supplemented with 5 g/L yeast extract and 0.1% L-cysteine (BHIS). The SOD-deficient *E. coli* strain QC779 was obtained from the *E. coli* Genetic Stock Center at Yale University. *E. coli* strains were cultured in LB at 37 °C.

### Generation of *C. difficile* SOR mutants

*C. difficile sor*::*erm* was generated in the BI-1 background by insertional disruption using the ClosTron method as previously described (34). Successful disruption was verified by gene-specific PCR (forward primer: CTCAGCTAACGGAACATGGCT, reverse primer: GGATCCAAATATTACAGTTCAGCCTTCC), the ClosTron ErmRAM PCR, and Southern blotting (34) (data not shown).

### Generation of complemented strains

Complemented strains were generated for the BI-1 background using the pMTL82153 backbone designed for constitutive expression in clostridia (64). The *sor* open reading frame and the 12bp upstream (containing the putative ribosome binding site) were amplified from *C. difficile* BI-1 genomic DNA by PCR (forward primer: CCATGGGAGGTGAGTTTATGTGTAGTGAAC, reverse primer: CTGCAGTTACAGTTCAGCCTTCCATAATC), TA-cloned into pGEM-T Easy (Promega, Madison WI, USA), and sub-cloned into pMTL82153 using *EcoR*I and *BamH*I restriction sites. Plasmids were transferred into *C. difficile* by conjugation as described by Heap et al. (65). Unless otherwise noted, “wild type” refers to *C. difficile* strain BI-1 harboring the empty pMTL82153, “*sor*::*erm*” is the BI-1 *sor* mutant bearing empty pMTL82153, and “*sor*::*erm*/p*sor*” is the BI-1 SOR mutant with the complementing plasmid.

### *C. difficile* aerotolerance assay

To measure *C. difficile* aerotolerance, liquid broth cultures were exposed to oxygen for defined time periods, and surviving bacteria were enumerated by plating. Standardized, mid-exponential phase cultures were prepared as follows: Cultures were grown to saturation under anaerobic conditions in BHIS, sub-cultured 1:50 in fresh BHIS, grown to mid-exponential phase (OD_600_=0.4-0.8, approximately 4-5 hours), and diluted in fresh BHIS to OD_600_=0.1. Cell numbers in the starting cultures were determined by plating. Cultures were removed from the anaerobic chamber and incubated in a 37°C shaker, with lids loosened to ensure aeration. Samples were removed at indicated time points, returned to the anaerobic chamber, diluted in pre-reduced BHIS broth, and plated on BHIS agar. Colony-forming units (CFU) of surviving bacteria were counted after 24 hours.

### Menadione and H_2_O_2_ sensitivity

*C. difficile* sensitivity to H_2_O_2_ and the redox-cycling compound menadione were measured by disc diffusion on TG agar. Standardized, mid-exponential cultures were prepared as for the aerotolerance assay. 100µL of culture was plated on TG agar and allowed to dry. Sterile filter paper discs impregnated with 125µg H_2_O_2_ (4µL of 3% solution) or 100µg MD (10µL of 10mg/mL solution, in EtOH) were placed on the dry plates. Plates were incubated anaerobically with no air exposure, or removed from the anaerobic chamber (15 min after discs were placed, to allow the H_2_O_2_ or MD to diffuse out into the agar) and incubated aerobically at 37°C for 15min. The plates were returned to the chamber and incubated anaerobically. Zones of clearance around the discs were measured after 24 hours.

### Expression of *C. difficile* SOR in SOD-deficient *E. coli*

As an independent approach to studying its function, *C. difficile sor* was cloned and expressed in QC779, an *E. coli* strain lacking native cytoplasmic superoxide dismutases. The *sor* open reading frame from *C. difficile* was PCR-amplified (forward primer: ATGTGTAGTGAACAAAAATTTTTTATATGT, reverse primer: CAGTTCAGCCTTCCATAATCCATG) and TA cloned into the expression plasmid pTrcHis2 TOPO (Thermofisher, Carlsbad CA, USA), in-frame with the C-terminal *myc*-6XHis and Myc tags, according to the manufacturer’s instructions. The resulting plasmid (p*sor*), or the empty vector, was transferred into QC779 with the quick transformation protocol of Chung et al (66). The strains are denoted as QC779 (WT), QC779/p*CDsor* (SOR-expressing) and QC779/vector (empty vector).

Cultures of QC779 and its derivatives (OD_600_=0.4) were treated with varying concentrations of IPTG for 2 hours at 37°C. The bacteria were then lysed by sonication (Heat Systems Ultrasonic Processor XL, 20% power, 3-10 10s pulses), separated on 4-20% gradient acrylamide gels (BioRad, Hercules CA, USA) and transferred to nitrocellulose. SOR expression was verified by immunoblotting against the Myc tag. Anti-Myc antibody (Protein Mods, Madison WI, USA) was used at 1:5,000 and the detecting anti-mouse antibody (Sigma-Aldrich, St. Louis MO, USA) at 1:8,000.

### ROS Sensitivity in *E. coli*

ROS sensitivity of the *C. difficile* SOR-expressing *E. coli* strain was measured as the zone of inhibition to menadione. Cultures were grown to mid-exponential phase in LB broth with 50μg/mL ampicillin, standardized to an OD of 0.1, plated (100µL) on LB agar and allowed to dry. Sterile filter paper discs impregnated with 400µg MD (40µL of 10mg/mL solution, in EtOH) were placed on the dry plates, which were incubated aerobically at 37°C. Zones of clearance around the discs were measured after 24 hours.

### In-gel assay for superoxide scavenging activity

The ability of *C. difficile* SOR to scavenge superoxide was tested by the in-gel method of Beauchamp et al. (37), following recommendations from Weydert et al. (38). Briefly, *E. coli* QC779 and its derivatives were grown to 0.8 OD (100mL in LB with 50µg/mL ampicillin and 25mM glucose), pelleted, and resuspended in LB with 20µM IPTG to induce SOR expression. After 2h, the bacteria were pelleted, resuspended in phosphate buffer (50µM phosphate, pH 7.8) and lysed by gentle sonication (Heat Systems Ultrasonic Processor XL, 20% power, 3-10 10s pulses). Cell debris was pelleted by centrifugation. Total protein in the supernatant (extract) was quantified with the Pierce 660 kit (Thermo Scientific, Waltham MA, USA). Extracts (20µg) were separated on a 12% Tris-HCl native gel (BioRad, Hercules CA, USA). Purified bovine superoxide dismutase (10µg, Sigma-Aldrich, St. Louis MO, USA) was used as a positive control (data not shown). After electrophoresis, the gel was soaked in a mixture of 2mg/mL nitroblue tetrazolium (NBT), 4.25µL/mL TEMED and 28µM riboflavin, and placed under a fluorescent light until clear bands appeared. Immunoblotting of QC779 and QC779/p*CDsor* lysates on the native gel was performed in parallel, using 1° mouse α-His at 1:1,000, and 2° goat α-mouse at 1:8,000.

### Xanthine oxidase assay for superoxide scavenging activity

To quantify the superoxide scavenging of the SOR-expressing *E. coli* QC779 strains, activity was measured using the Superoxide Dismutase Colorimetric Activity Kit (Invitrogen), according to manufacturer’s instructions, with modifications. Briefly, bacterial lysates were prepared as for the in-gel assay. 50µg of the extracts were diluted 1:1 (v/v) with Assay Buffer, then incubated with 1X xanthine oxidase and 1X substrate at room temperature. Absorbance was read at 450 nm at multiple time points. ROS scavenging activity was measured as 1/Abs450.

### Antibiotic sensitivity

Sensitivity to metronidazole, rifampicin, cefoxitin sodium, and trimethoprim was assessed as the minimum inhibitory concentration (MIC) by broth microdilution. Sensitivity to penicillin, streptomycin, kanamycin, chloramphenicol, erythromycin, and moxifloxacin was measured as the zone of inhibition using commercially prepared discs. In all cases, mid-log cultures standardized to 0.1OD (prepared as for the aerotolerance assay) were used. For MICs, 2-fold serial dilutions of antibiotics were made in 96-well plates and inoculated 1:10 with standardized mid-exponential cultures. For zones of inhibition (ZOIs), 100uL of the standardized mid-log culture were plated on Wilkins-Chalgren agar. For the air-exposed condition, 96-well plates were shaken aerobically at room temperature for 20 minutes and agar plates for ZOIs were exposed for 10 minutes before incubating anaerobically. Both MICs and ZOIs were determined after 24h.

### Spore and toxin production

To measure spore production, 12-day old liquid cultures were pelleted, washed in water four times, heat shocked at 65° for 15 minutes, and plated on Brain Heart Infusion (BHI) agar supplemented with taurocholate. For a qualitative assessment, the spore preparations were also examined by phase microscopy. Toxin production was measured with the Wampole *C. difficile* Toxin A/B ELISA kit (Alere, Waltham MA, USA), using supernatants from 3-day old broth cultures. Toxin measurements were normalized to total protein amount in the supernatant.

### Aerotolerance assay protein and peptide processing for LC-MS/MS analysis

Samples (10 ml) of 4 strain-treatment combinations from the *C. difficile* aerotolerance assay (WT and *sor::erm* derivatives, in anaerobic conditions and with 30 minutes of oxygen exposure) were collected and centrifuged at 3200 x g, 4°C for 10 minutes. Cell pellets were then resuspended in 50 mM ammonium bicarbonate and lysed via 6 30-second sonication on ice. Cell lysates were centrifuged at 16,000 g, 4°C for 30 minutes and protein concentration of the supernatants were determined using BCA assay. Trypsin digestion was performed on 10 µg of protein following standard Proteasemax-assisted digestion protocol. Pierce C18 spin columns were used to clean up peptide digests.

### LC-MS/MS run parameters

Nano liquid chromatography–tandem mass spectrometry (LC–MS/MS) analysis was carried out using an online reversed phase nano-LC Agilent 1200 coupled with LTQ-Orbitrap Velos mass spectrometer. Peptide digests (3 µl) in 3% acetonitrile 0.1% formic acid were injected into a C18 PepMap trap column (75 µm ID × 2 cm, Dionex) and separation was achieved on 5–95% acetonitrile gradient (120 min) at a flow rate of 0.4 μL/min through a 15 cm long C-18 picofrit column (100 μm internal diameter, 5 μm bead size, Nikkyo Technos Co., Tokyo, Japan) installed on the online nano electrospray ionization (NSI) source of the mass spectrometer. Ionization was performed using an electrospray voltage of 2 kV and ions were then channeled into an LTQ Orbitrap Velos operated in the positive mode to record full scan MS spectra (m/z 300–2000) at a resolution of 30,000 followed by isolation and fragmentation of the 5 most intense ions by collision-induced dissociation.

### Label-Free quantitative proteomic analysis

Raw MS data was processed with MaxQuant software (version 1.2.0.18), and identification and quantification of proteins from spectral data was carried out using MaxQuant’s label-free quantification algorithm searching *C. difficile* BI-1 (GCF_000211235.1_ASM21123v1) and contaminant database appended with reverse sequence decoy database. Scaffold 5.1.2 was used to compile and analyze the label-free quantification (LFQ) intensities of the proteins from the MaxQuant analysis. The false discovery rate (FDR) for peptide identification was set to 5% and fold changes greater than 2 was arbitrarily set as the cutoff defining differentially regulated proteins.

### Functional classification, gene set enrichment and network analysis

Functional classification analysis of differentially regulated genes from 4 of the 12 possible ratios (Table 1) were performed using hypergeometric testing (BiNGO) and gene set enrichment analysis (GSEA). Overrepresented gene classifications and processes from gene ontology (GO), Kyoto Encyclopedia of Genes and Genomes (KEGG), and Clusters of Orthologous Genes (COG), were used as guide in constructing STRING database v.11.5 protein-protein interaction (PPI) networks visualized in Cytoscape v. 3.9.1.

### Statistical Analysis

The Excel-Stat application was used for statistical analysis. Student’s t-tests were used to compute differences between two groups, and errors bars calculated from standard deviation(s). Analysis of Variance (ANOVA) was used to compare >2 groups, followed by Tukey’s HSD test for post-hoc analysis.

## Acknowledgements

We thank Keith Lammers for expert technical assistance. This work was supported by funding from the National Institutes of Health [R21 AI119478 (GV) and the US Dept. of Veterans Affairs [IK6BX003789(GV; Research Career Scientist Award), and I01BX001183(GV)].

## Data Availability Statement

The datasets presented in this study can be found in online repositories. The names of the repository/repositories and accession number(s) can be found in the article/Supplementary Material.

## Competing Interests Statement

The authors declare no competing interests.

